# Lethal mutagenesis of Rift Valley fever virus induced by favipiravir

**DOI:** 10.1101/601344

**Authors:** Belén Borrego, Ana I. de Ávila, Esteban Domingo, Alejandro Brun

## Abstract

Rift Valley fever virus (RVFV) is an emerging, mosquito-borne, zoonotic pathogen with recurrent outbreaks paying a considerable toll of human deaths in many African countries, for which no effective treatment is available. In cell culture studies and with laboratory animal models, the nucleoside analogue favipiravir (T-705) has demonstrated great potential for the treatment of several seasonal, chronic and emerging RNA virus infections of humans, suggesting applicability to control some viral outbreaks. Treatment with favipiravir was shown to reduce the infectivity of Rift Valley fever virus both in cell cultures and in experimental animal models, but the mechanism of this protective effect is not understood. In this work we show that favipiravir at concentrations well below the toxicity threshold estimated for cells is able to extinguish RVFV from infected cell cultures. Nucleotide sequence analysis has documented RVFV mutagenesis associated with virus extinction, with a significant increase in G to A and C to U transition frequencies, and a decrease of specific infectivity, hallmarks of lethal mutagenesis.

## INTRODUCTION

Rift Valley fever virus (RVFV) is a mosquito-borne bunyavirus belonging to the recently reclassified phlebovirus genus (Family *Phenuiviridae*, Order Bunyavirales). RVFV causes an important disease in ruminants often transmitted to humans after the occurrence of epizootic outbreaks. Although the disease has been reported only in African countries with some incursions in the Middle East and Indian Ocean islands, there are concerns for its potential spread to other locations, including Europe (1, 2). Currently, there is no available treatment or licensed RVF vaccine in Europe, therefore the development of effective control strategies is an essential field of research. The use of antiviral agents for treatment in livestock is generally not affordable, although it is often being proposed for viral veterinary diseases as a strategy to fill the gap between the time of the infection and the effective development of the host immune response (3, 4). In contrast, in the case of humans infected or at risk of infection with RVFV, antiviral treatment is fully warranted. Antiviral base and nucleoside analogues effective against RVFV are available (5-7). Among the nucleoside analogs exerting anti-RVFV activity, ribavirin [1-β-D-ribofuranosyl-1-H-1,2,4-triazole-3-carboxamide], favipiravir [6-fluoro-3-hydroxy-2-pyrazinecarboxamide] and more recently BCX4430 [(2S,3S,4R,5R)-2-(4-amino-5H-pyrrolo[3,2-d]pyrimidin-7-yl)-5-(hydroxymethyl)pyrrolidine-3,4-diol] (Galidesivir) have been described (6-8). Particularly, favipiravir displays potent antiviral activity against different RNA viruses (9-16). Upon cell intake, favipiravir is converted by cellular enzymes into its active form (favipiravir-4-ribofuranosyl-5-triphosphate), which functions as a purine nucleotide analogue. Favipiravir can act as an RNA chain terminator (17, 18), and as lethal mutagen for several RNA viruses, including hepatitis C virus (19), foot-and-mouth disease virus (20), West Nile fever virus (21), norovirus (22), and influenza virus (23), among others. Here we show that the mechanism of action of favipiravir against RVFV in cell culture is due, at least in part, to the accumulation of mutations in the viral genome that leads to a progressive decrease in viable viral progeny. This conclusion is based on a significant increase of mutation frequency measured in a 1060 bp fragment from the Gc (glycoprotein C)-coding region, and a concomitant decrease of specific infectivity, associated with virus extinction, supporting the use of favipiravir for the control of emerging RVFV infections in humans.

## RESULTS

### Effect of favipiravir (T-705) on RVFV 56/74 progeny virus yield

The aim of this work was to determine whether favipiravir induces lethal mutagenesis upon RVFV infection in cell culture. We first determined the effect of different concentrations of favipiravir on the yield of RVFV isolate 56/74 in Vero cells. The half inhibitory concentration (IC_50_, concentration required for a 50% decrease in the production of infectious progeny), was calculated to be 6.22 ± 1.5 μM (average of 3 determinations) **(Figure 1).** Since the cytotoxic concentration 50 (CC_50_) of T-705 for Vero cells is >10000 μM (21), the therapeutic index for RVFV (TI = CC_50_ / IC_50_) is > 1600.

**Figure 1.**
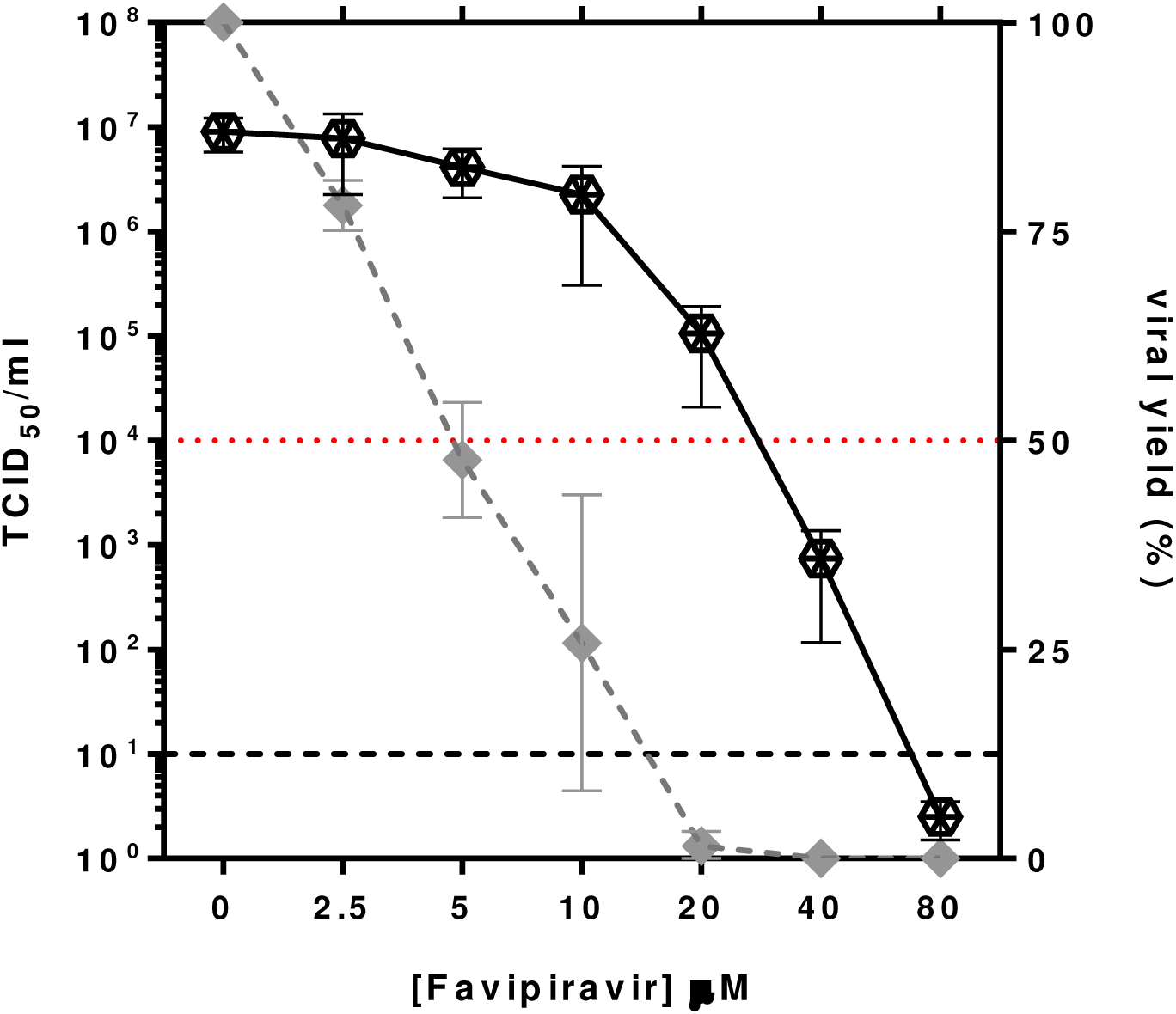
Effect of favipiravir treatment on viral yields in cell culture. Vero cells were pretreated overnight with the indicated concentrations of drug and then infected with RVFV 56/74 at a moi of 0.1 PFU/cell. Infection proceeded for 72 hpi in the presence of the same concentration of favipiravir. Results are shown either as TCID_50_/ml (continuous line; left Y axis) or as percentages of RVFV titer with respect to the one obtained in untreated cells (dashed line; right Y axis). Values under the limit of sensitivity of the assay (estimated to be 10, indicated with a grid line) were arbitrarily represented as 2. Red line indicates 50%. Error bars denote SD.

The RVFV 56/74 isolate was subjected to serial passages in Vero cells in the absence or presence of different concentrations of favipiravir (**Figure 2a**). The titration of supernatants indicated loss of infectivity in the passages carried out in the presence of 80 μM favipiravir, as well as for higher doses, such as 160, 200 and 500 μM, but not at the lowest concentrations tested. A concentration of 20 μM produced a slight reduction in viral yield during the consecutive passages, whilst at 40 μM the progeny production declined progressively, and was not detected after the fifth, sixth and seventh passage. Interestingly, virus yields were recovered at passages eighth and ninth. These data suggested the selection of a resistant mutant under certain suboptimal inhibitory concentrations of the drug. To test this hypothesis, viral yields in the presence of different concentrations of favipiravir were analyzed for this virus. As expected from the previous results (**Figure 1**), favipiravir concentrations over 5 μM significantly reduced the yield of either the parental or the serially passaged virus RVFV 56/74 not treated with the drug, whilst for the virus selected after 8 passages in the presence of 40 μM favipiravir a 50% reduction in viral yield was achieved at a higher concentration, rendering an IC_50_ of 19.2 μM (**Figure 2b**). These results clearly indicated a 3-fold change in susceptibility to the drug when compared to the parental strain, although it remained susceptible to higher concentrations of the drug. The characterization of this resistant variant is currently under investigation.

**Figure 2.**
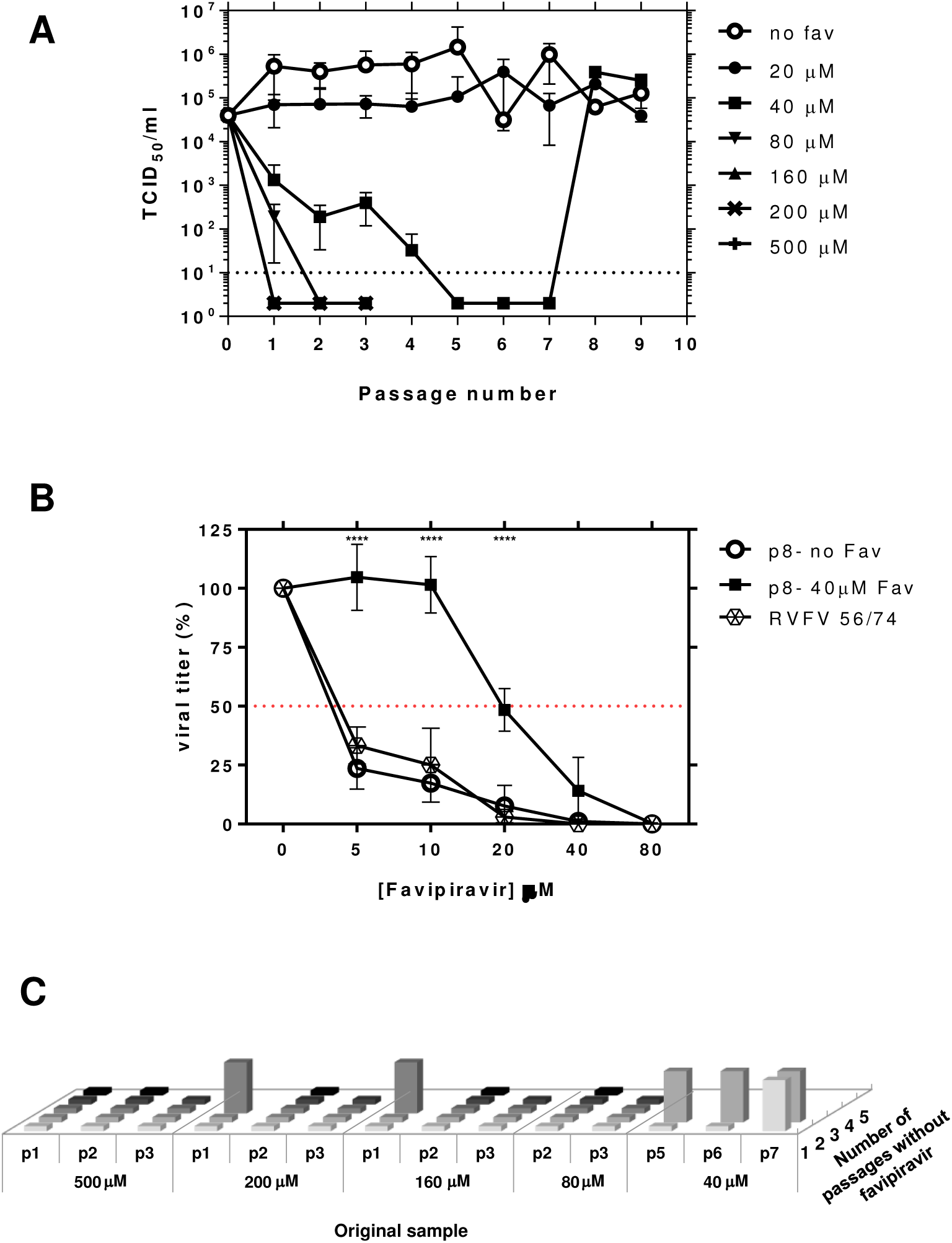
Serial passages of RVFV in Vero cells in the absence or presence of favipiravir. A) Viral titers of RVFV after each passage in the absence or presence of different concentrations of favipiravir (T-705), as indicated. For concentrations higher than 80 μM, only 3 passages were performed. Viral titers in the supernatant of infected cells were determined at 72 hpi. Error bars denote SD. B) Comparison of the viral titers obtained from virus recovered after passage 8^th^ (previously propagated or no with 40μM favipiravir) and parental RVFV56/74 in the presence of a 5-80 μM range of favipiravir concentrations. Titers are represented as percentages with respect to those obtained without drug. C) Lack of detectable infectivity after blind passages as a criterion for RVFV extinction. Cell culture supernatants from passages carried out in the presence of favipiravir where no cytopathic effect (cpe) was observed (Figure 2A, samples whose titres were below the limit of sensitivity of the assay) were subjected to additional passages (up to five) in the absence of drug. Original samples are named according to their passage number in the presence of the indicated favipiravir concentration (in μM). Flattened columns indicate no cpe detection; lifted columns indicate evident cpe.

To confirm virus extinction, cell culture supernatants from passages carried out in the presence of favipiravir where no cytopathic effect (cpe) had been observed were subjected to up to five additional passages in the absence of the drug (**Figure 2c**). No infectivity was rescued from samples that were collected after at least two serial passages in the presence of 80 μM favipiravir or higher. Viral infectivity could be recovered from supernatants of cells previously treated with 40 μM favipiravir. These data confirmed the previous observation, indicating that the deleterious effect on virus propagation at this concentration was not complete.

### Analysis of RVFV passaged in absence or presence of favipiravir

Decrease of specific infectivity, broadening of the mutant spectrum and invariance of the consensus sequence of the viral population are diagnostic of lethal mutagenesis (19-21, 23-28). To explore these parameters, RVFV populations at passages 1 to 4 in absence or presence of 40 μM favipiravir were examined. Extracellular viral titers and viral RNA decreased significantly in the presence of favipiravir (**Figure 3**). The changes in infectivity and viral RNA production resulted in a gradual decrease of specific infectivity in the populations passaged in the presence of favipiravir (up to 8-fold in four passages) but not in its absence (p<0.0001 for passages 1 to 4; t test). Thus, the first diagnosis criterium of lethal mutagenesis is fulfilled for favipiravir acting on RVFV.

**Figure 3.**
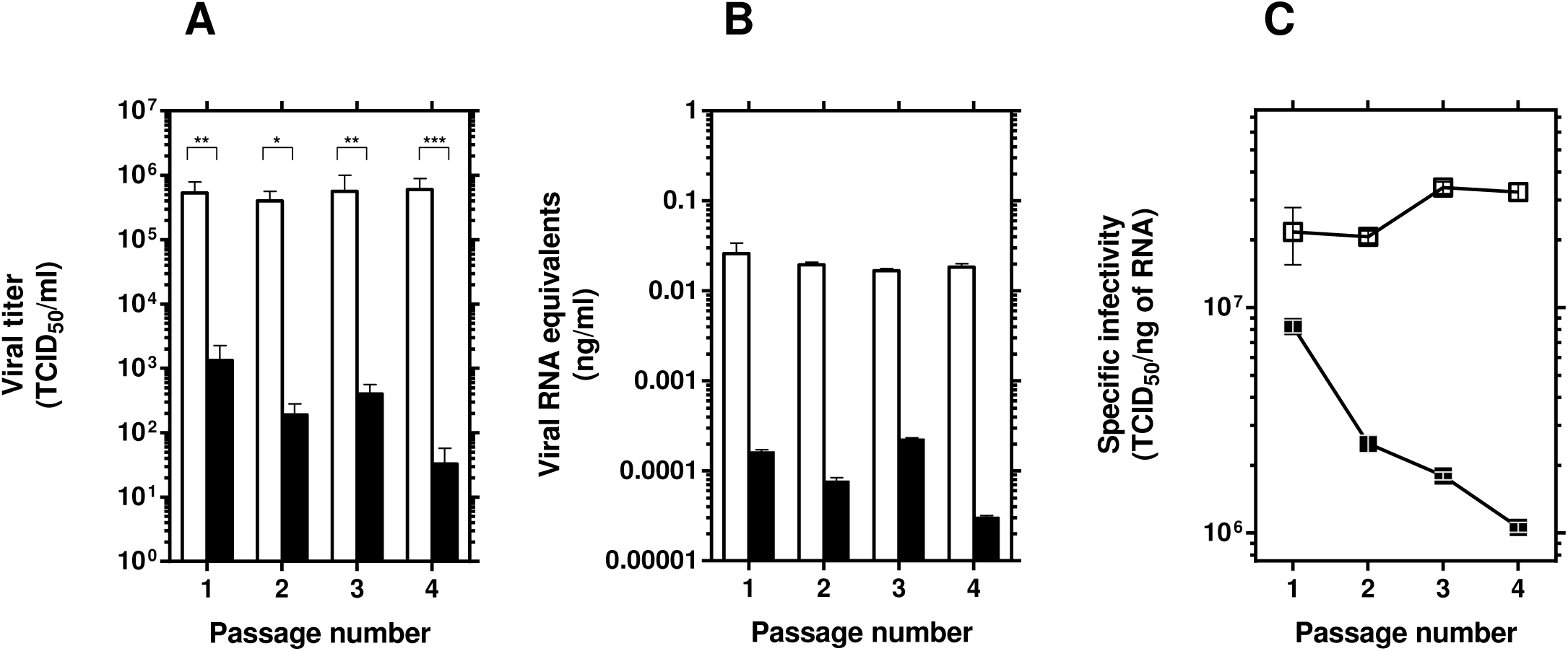
Effect of favipiravir in the specific activity of RVFV. A) Viral titers of RVFV after each passage (n=4) in the absence (white columns) or presence (black columns) of 40 μM favipiravir (redrawn from data shown in Figure 2A). B) Quantification by qRT-PCR (in triplicate) of RNA extracted from the supernatants of untreated (white columns) or 40 μM favipiravir-treated (black columns) Vero cells after each passage (n=4). C) Specific infectivity upon passage of RVFV in the absence (open symbols) or presence (closed symbols) of 40 μM favipiravir. Values correspond to the ratio between infectivity (panel A) and the amount of viral RNA (panel B). Error bars denote SD. Statistically significant differences are indicated by asterisks (*p≤0.05, **p≤0.01, ***p≤0.005, ****p≤0.0001; Student’s t-test).

To examine whether the decrease in progeny production was associated with increased mutagenesis of the viral RNA, total RNA from the cell culture supernatant of passage 4 in the absence or presence of 40 μM favipiravir was extracted and a 1.1kb DNA fragment encoding a glycoprotein Gc region comprising nucleotides 2091 to 3223 (numbering according to GenBank sequence of RVFV strain SA-75, accession number DQ380189) was amplified by RT-PCR and subjected to molecular cloning and nucleotide sequencing. For each condition (with/without drug) a total of 29 clones were analyzed, corresponding to 31,610 nucleotides sequenced. The results (**Table 1**) indicated a significant 2.5-fold increase of minimum mutation frequency associated with replication in the presence of favipiravir (p< 0.0001; χ2 test). Of the total 142 different mutations identified in the populations analyzed, 89 were detected only in the population passaged in the presence of favipiravir, 4 were detected only in the population passaged in the absence of favipiravir, and 49 were common in the two populations (indicated in **Table S1**). Once the common mutations are excluded, the minimum mutation frequencies are 1.3 × 10^−4^ and 2.8 × 10^−3^ in absence and presence of favipiravir, respectively, and the difference is statistically significant (P<0.0001; χ2 test). The distribution of the number of Gc sequences was analyzed as a function of the number of mutations per clone (**Figure 4**). While the viral population that was passaged in absence of drug yielded 7 clones without mutations and 7 clones with only one mutation (24.1% in both cases), only 1 clone without mutations and 2 with one mutation were retrieved from the population that was passaged in the presence of favipiravir (1/29, 3.4%; 2/29, 6.9%, respectively). Finally, the consensus genomic nucleotide sequence (analyzed at passage 4) did not change as a result of replication in the presence of favipiravir, an observation already found during lethal mutagenesis of other RNA viruses (19, 27-30).

**Table 1.** Mutant spectrum analysis by molecular cloning and Sanger sequencing of glycoprotein Gc coding region of RVFV passaged 4 times in Vero cells

| **RVFV**<br>**population<sup>a</sup>** | **Number of nucleotides**<br>**analysed**<br>**(clones/haplotypes)<sup>b</sup>** | **Different**<br>**(total)**<br>**mutations<sup>c</sup>** | **Minimum**<br>**mutation**<br>**frequency<sup>d</sup>** | **Maximum**<br>**mutation**<br>**frequency<sup>e</sup>** | **Different**<br>**synonymous**<br>**(non-synonymous)**<br>**mutations** | **Total**<br>**synonymous**<br>**(non-synonymous)**<br>**mutations** |
| --- | --- | --- | --- | --- | --- | --- |
| **56/74 p4, no**<br>**drug** | 31,610 (29/21) | 53 (375) | $1.7 \times 10^{-3}$ | $1.2 \times 10^{-2}$ | 48 (5) | 354 (21) |
| **56/74 p4 +**<br>**favipiravir** | 31,610 (29/29) | 138 (348) | $4.4 \times 10^{-3}$ | $1.1 \times 10^{-2}$ | 85 (53) | 284 (64) |

<sup>a</sup>The origin and passage history of the viral populations analysed is described in Fig. 2 and Materials and Methods.
<sup>b</sup>The genomic region analysed spans residues 2110-3199 of glycoprotein Gc coding region; the residue numbering is that of the SA-75 genome. The values in parenthesis indicate the number of clones analysed followed by the number of haplotypes (number of different RNA sequences).
<sup>c</sup>The number of different and total mutations were counted relative to the consensus sequence of the corresponding population. Different and total mutations were used to calculate the minimum and maximum mutation frequency, respectively.
<sup>d</sup>Data represent the average number of different mutations per nucleotide in the mutant spectrum relative to the consensus sequence of the corresponding population. The results indicate a significant 2.5-fold increase of minimum mutation frequency associated with replication in the presence of favipiravir ( $p < 0.0001$ ; $\chi^2$ test)
<sup>e</sup>Data represent the average number of total mutations per nucleotide in the mutant spectrum relative to the consensus sequence of the corresponding population.

**Figure 4.**
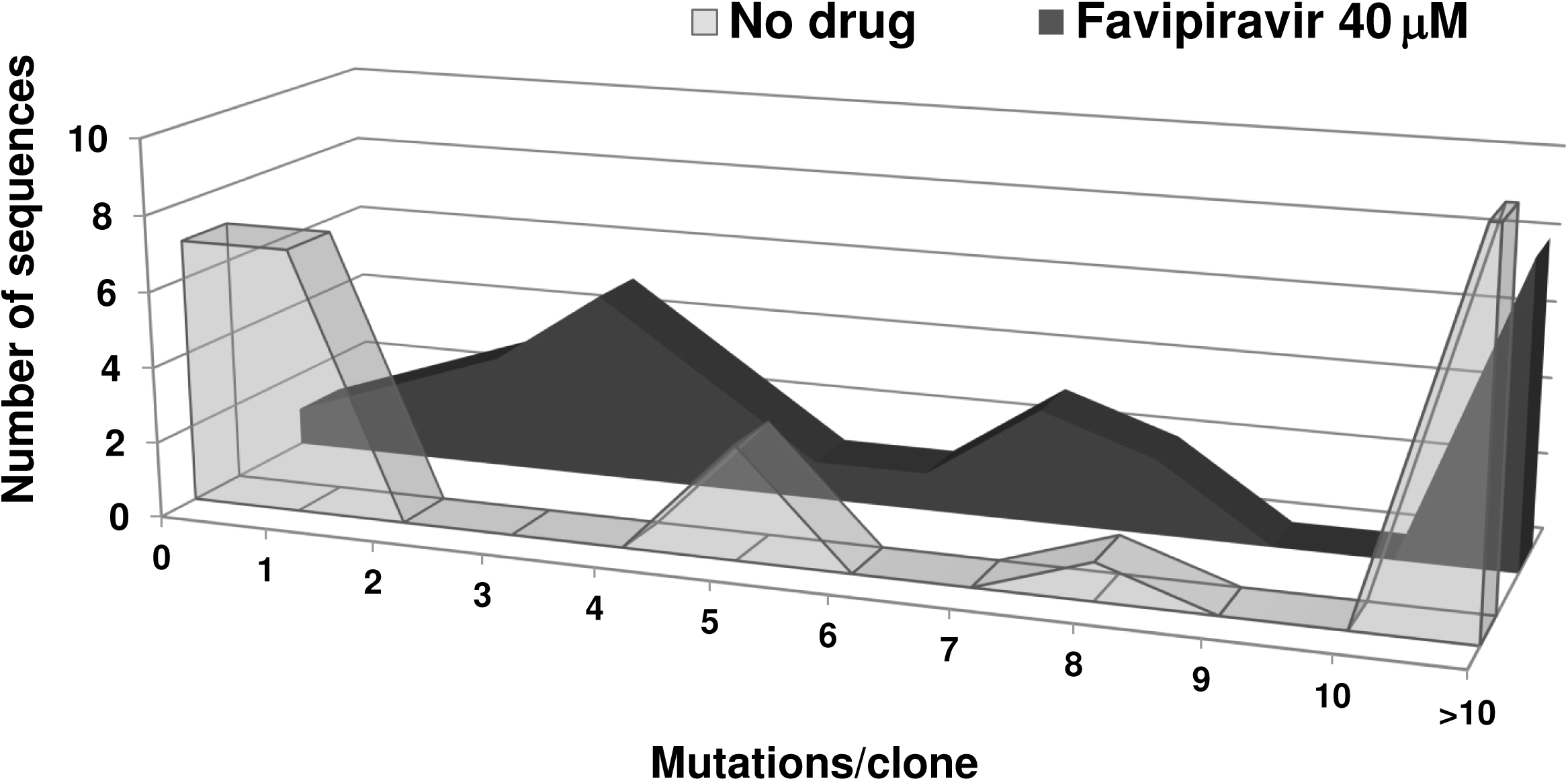
Distribution of mutations among individual clones. Plot of the distribution of the number of mutations per molecular clone in the RVFV populations passaged in the absence or presence of 40 μM favipiravir.

### Analysis of mutation types

In order to analyze whether passage of RVFV in the presence of favipiravir resulted in any deviation of mutation types, we compared the mutation matrices in the mutant spectrum of the populations passaged in absence or presence of the drug. The results indicate a predominance of G to A and C to U transitions associated with replication in the presence of favipiravir: without drug the number of changes G→A + C→U was 24 (10+14), while with favipiravir it was 98 (52+46), thus resulting in 3-fold increase of the ratio (G→A) + (C→U) / (A→G) + (U→C) (**Figure 5**). The difference between the type of mutations was statistically significant (p< 0.0001; χ2 test), and reveals that favipiravir produces a bias in favor of GA and CU transitions; the transition to transversion ratio was comparable: 12.2 and 18.7 upon passage in absence and presence of favipiravir, respectively.

**Figure 5.**
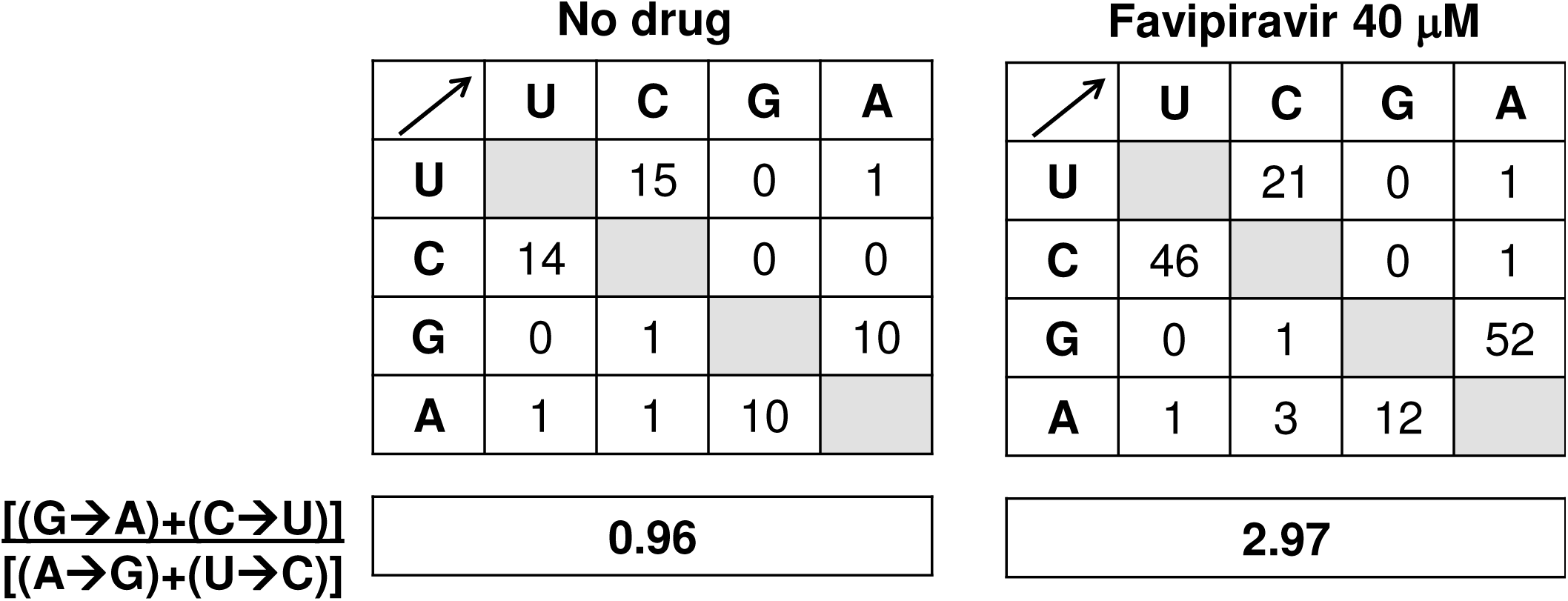
Mutational spectrum induced by favipiravir on RVFV. Matrices of mutation types found in the glycoprotein Gc coding region of RVFV passaged 4 times in Vero cells in the absence or presence of 40 μM favipiravir. The boxes below each matrix quantify the mutational bias, according to the ratio shown on the left.

These results indicate that favipiravir is a mutagenic agent for RVFV and suggest that its anti-RVFV activity may be exerted at least in part through lethal mutagenesis.

## DISCUSSION

The broad spectrum antiviral activity of favipiravir has been established with representatives of several families of RNA viruses, so it stands as a promising drug to be used against viral pathogens. In the case of Rift Valley fever, for which no therapeutic or prophylactic treatments are available for humans, favipiravir could be a powerful therapeutic tool to provide immediate treatment to farmers, veterinary and research personnel at risk of infection after being in contact with the virus or virus-infected animals. Favipiravir was initially considered an inhibitor of viral RNA synthesis, exerted mainly through RNA chain termination (17, 18). However, model studies with several viruses in cell culture and *in vivo* suggest that the antiviral activity of favipiravir may include lethal mutagenesis.

Previous studies have shown that favipiravir treatment of laboratory animals exposed to RVFV increased their survival, and counteracted acute or neurological disease syndromes (6, 7). Furthermore, treatment efficacy was enhanced when favipiravir was combined with ribavirin treatment (7). Knowledge of the mechanism of action of this promising drug is important to improve antiviral designs. Our data clearly show at least a 2.5-fold increased RVFV mutation frequencies when the virus was propagated in the presence of 40 μM favipiravir, a dose which is 10-to 25-fold lower than the dose proven effective for other RNA viruses tested in a similar way (19-21). The limitation of non-synonymous mutations introduced during the four passages in the absence of favipiravir (9.4% and 5.6% of the total 53 and 375 for different and total mutations, respectively) is diminished in the passages in the presence of favipiravir (38.4% and 18.4%, respectively), consistent with random introduction of mutations that decrease viral fitness.

One of the most advantageous features of favipiravir is that it shows a high barrier for the generation of resistance mutants (9). Since its activity against human influenza virus was first described (11), only one resistant mutant has been reported (31). Attempts to isolate resistant variants of polio-and influenza viruses seem to have been unsuccessful (32-34). In addition, influenza virus pairs isolated from patients before and after favipiravir administration showed no changes in the susceptibility to the drug (35). Similar results were reported in norovirus-infected mice after favipiravir treatment (22). Moreover, previous studies reporting favipiravir mediated viral extinction by lethal mutagenesis did not describe selection of specific mutations or resistant variants (19-21, 23). In our system, 40 μM favipiravir seemed not to lead to the extinction of the virus, since after a lag phase of three consecutive blind passages the presence of cpe was detected again in cultures. The comparison of the IC_50_ value of the virus recovered in supernatant after the lag phase with that of the original samples confirmed the selection of variants with increased favipiravir resistance, suggesting the selection of mutant viruses with increased resistance to the drug. Work is currently underway in order to characterize this novel resistance variant both at genomic and phenotypic level *in vitro* and *in vivo*.

The appearance of variants showing decreased susceptibility to the drug has been shown earlier (Delang et al., 2014) and it highlights the importance of deciphering the exact mechanisms exerted by new promising antiviral drugs, such as favipiravir. In this sense, drug concentration could be proposed as a crucial determinant on the antiviral effect. Thus, between high concentrations -leading to a total abolishment of infectivity- and low concentrations -with none or low effects on viral replication-there might be a critical intermediate concentration that, even though forcing the virus into lethal mutagenesis, promotes the selection for drug resistant variants before the virus is extinguished. Nonetheless, it is worth noting that other studies found that drug concentrations close to the IC_50_ led to viral extinction of influenza virus without the emergence of variants (23) perhaps indicating that other unknown factors play a role the inhibitory mechanism of T-705. Still, the antiviral activity mechanism of favipiravir is not fully understood. Most nucleoside analogues used as antiviral agents display several mechanisms of activity (36-38). This is also the case for favipiravir with a dual chain terminating and mutagenic activity by ambiguous base pairing, as well as a possible role in modulation of innate immunity (10).

In summary, what our results suggest is that favipiravir can act as a lethal mutagen for RVFV but they do not exclude that other mechanisms, alone or together with lethal mutagenesis, are required for virus extinction.

## MATERIALS AND METHODS

### Cells, viruses and infections

Vero cells (ATCC Cat.No. CCL-81) were grown in Dulbecco’s modified Eagle’s medium supplemented with 5% –10% fetal calf serum (FCS), and L-glutamine (2 mM), penicillin (100 U/ml) and streptomycin (100 μg/ml), in a humid atmosphere of 5% CO_2_ at 37°C.

The virus used in this study originates from a sheep experimentally infected in our laboratory (39) with RVFV isolate 56/74 (40, 41). The virus was re-isolated from infected sheep plasma by culturing in a c6/36 mosquito cell line (ATCC Cat.No. CRL 1660). The infection of cells in the absence of drugs and the assays for titration of infectivity in liquid medium to determine tissue culture infective doses (TCID) were performed as previously described (42). Titers were determined according to (43). Assays to quantify plaque-forming units (pfu) were done in semisolid medium including 1% Carboxymethylcellulose (CMC, Sigma). Monolayers were fixed and stained 3-5 days post infection.

### Infection in the presence of favipiravir

Favipiravir T-705 (Atomax chemicals Co. Ltd., China) was dissolved in DMSO at a concentration of 100mM, aliquoted, and stored at -70°C. For use in the experiments, the solution was thawed and diluted in DMEM to reach the desired concentration indicated in each experiment. Cells were pretreated with favipiravir 14 h prior to infection, the virus was adsorbed for 1 h to cells in absence of drug, and then infection continued in the presence of the same drug concentration.

Serial passages in the presence of favipiravir were performed by infecting Vero cell monolayers seeded on M6 plates, in 2 ml of cell culture medium; 0.5 ml of the supernatant was collected at 72 h post-infection, and used for the next infection. The first passage was carried out at a multiplicity of infection (MOI) of 0.1 PFU/cell (4 ×10^5^ cells infected with 4 ×10^4^ PFU of virus). Mock-infected cells were maintained in parallel to control for the absence of contaminating virus. When no cytopathic effect (cpe) was observed, after several passages in the presence of favipiravir, the cell culture supernatant was used to perform up to 5 blind passages in the absence of drug, to ascertain that no infectious virus could be recovered. Lack of detectable cpe before and after blind passages was considered as criterion of RVFV extinction. The IC_50_ of T-705 was calculated as described previously (19).

### RNA extraction, RT-PCR, and nucleotide sequencing

RNA was extracted from the supernatants of infected cells using the Speedtools RNA virus extraction kit (Biotools B&M Labs, S.A., Madrid, Spain) according to the manufacturer’s instructions. RT-PCR was performed using SuperScript IV reverse transcriptase and Phusion High-Fidelity DNA polymerase (Thermo Scientific). cDNA was obtained by retrotranscription from RVFV M-segment RNA using primer MRV1ag (5’-CAAATGACTACCAGTCAGC-3’; positions 772-790; antigenomic orientation) as described (44). The cDNA was then amplified by PCR using oligonucleotides startGc forward (5’-ACTCGCATTGTCGACGTTTT) (positions 2091-2109; genomic orientation) and sm3 reverse (5’-GGATTAAGGAAGCGGGAAAAGCCC) (positions 3200-3223; antigenomic orientation). The amplified cDNA is an 1100 nt fragment that corresponds to the amino-terminal half of glycoprotein Gc, and it was chosen because it is highly conserved in RVFV (45). Amplifications in the absence of RNA were carried out in parallel to ascertain the absence of contaminating templates. For the analysis of mutant spectra, amplifications were carried out with template preparations diluted 1:10, 1:100 and 1:1000. When amplification of the 1:100 diluted template yielded a detectable DNA fragment, cloning was performed with the amplified fragment obtained with the undiluted template, as described previously (46). This procedure ensures an excess of template viral RNA molecules thus avoiding redundant amplifications of the same initial RNA molecules. Amplified DNA fragments were then ligated to pJET2.1 plasmid (Thermo Scientific), and the products transformed into *E.coli* DH5α. The pJET2.1 vector contains the lethal gene eco47IR enabling positive selection of recombinant plasmids. Plasmid DNA from individual bacterial colonies was purified and subjected to nucleotide sequencing, as previously described (47).

### Quantification of RVFV RNA

For the calculation of specific infectivity (the ratio of infectivity to viral RNA in a virus sample) a real time quantitative one-step PCR (RT-qPCR) of a fragment within L-segment RNA was carried out using RNA HighScript Master mix (Biotools B&M), according to manufacturer’s instructions. Primers used were L-forward (5’-TTCTTTGCTTCTGATACCCTCTG-3’) [5’ position 2872; sense orientation] and L-reverse (5’-GTTCCACTTCCTTGCATCATCTG) [5’ position 3006; antisense orientation] that yield a 135bp fragment within the RdRp-coding region (41). Detection was achieved by a L-segment specific hydrolysis probe (5’*-*^[6-FAM]^TTGCACAAGTCCACACAGGCCCCT[BHQ^1]^-3’). Quantification of RNA was relative to a standard curve obtained with known amounts of RVFV RNA, synthesized by *in vitro* transcription of a NdeI-linearized pGEM-T plasmid containing the 135bp cDNA target fragment using the T7 RiboMAX in vitro transcription kit (Promega). Negative controls (without template RNA and RNA from mock-infected cells) were run in parallel with each amplification reaction, to ascertain absence of contamination with undesired templates.

## Statistical analysis

Data analysis was performed using GraphPad Prism software version 6.

## Acknowledgements

We thank Nuria de la Losa for excellent technical assistance. This work was supported by grants S2013/ABI-2906 (PLATESA), P2018/BAA-4370 (PLATESA2) from Comunidad de Madrid/FEDER. AGL2014-57430-R and AGL2017-83326-R from Ministerio de Economía y Competitividad. SAF2014-52400-R and SAF2017-87846-R from Ministerio de Ciencia, Innovación y Universidades. CIBERehd (Centro de Investigación en Red de Enfermedades Hepáticas y Digestivas) is funded by Instituto de Salud Carlos III. Institutional grants from the Fundación Ramón Areces and Banco Santander to the CBMSO are also acknowledged.

