## Supplemental table S1 for "Lethal mutagenesis of Rift Valley fever virus induced by favipiravir"

| **Table S1. Mutations, corresponding amino acid and point accepted mutation (PAM) of the glycoprotein Gc coding region in the mutant spectra RVFV subjected to four passages in the absence or presence of 40 µM Favipiravir (T-705)** | | | | | |
|---|---|---|---|---|---|
| **No drug** | | | **40 µM favipiravir** | | |
| **Mutation[a]** | **Amino acid substitution** | **PAM250[b]** | **Mutation[a]** | **Amino acid substitution** | **PAM250[b]** |
| G2073A (8) | - | - | **G2045A** | S698N | 1 |
| T2079C (8) | - | - | **C2046T** | - | |
| A2094G (7) | - | - | **C2055T** | - | |
| G2112A (7) | - | - | **A2056C** | T702P | 0 |
| A2148G (8) | - | - | **C2057T** | T702I | 0 |
| G2154A (8) | - | - | **C2058T** | - | |
| T2172C (8) | - | - | **C2060T** | T703I | 0 |
| A2181G (8) | - | - | **G2063A** | C704Y | 0 |
| C2184T (8) | - | - | **C2064T** | - | |
| C2259T (9) | - | - | **C2066T** | S705F | -3 |
| **C2283T** | - | - | G2073A (4) | - | |
| T2292C (9) | - | - | **G2077A** | V709I | 4 |
| A2295G (9) | - | - | T2079C (4) | - | |
| G2328A (9) | - | - | **C2082T** | - | |
| T2340C (8) | - | - | **A2083C** | T711P | 0 |
| G2349A (8) | - | - | **G2088A** | - | |
| A2382G (8) | - | - | A2094G (4) | - | |
| T2388C (8) | - | - | **G2097A** | - | |
| G2418A (7) | - | - | **C2103T** | - | |
| C2427T (6) | - | - | **C2105T** | T718I | 0 |
| C2466T (6) | - | - | **T2111C** | L720S | -3 |
| T2478C (6) | - | - | G2112A (5) | - | |
| T2502C (6) | - | - | **G2131A** | G727R | -3 |
| C2514T (6) | - | - | **G2133A** | - | |
| G2531C (6) | S860T | 1 | **C2141T** | A730V | 0 |
| C2541T (6) | - | - | **T2145C** | - | |
| A2544T (6) | - | - | A2148G (4) | - | |
| T2575C (7) | - | - | G2154A (5) | - | |
| T2640C (8) | - | - | **G2160A** | - | |
| T2655C (8) | - | - | T2172C (3) | - | |
| T2664C (7) | - | - | A2181G (3) | - | |
| T2754C (8) | - | - | C2184T (3) | - | |
| **C2759T** (3) | S936L | -3 | **G2219A** | C756Y | 0 |
| G2763A (6) | - | - | **G2223A** | - | |
| C2766T (6) | - | - | **G2227A** | G759S | 1 |

| | | | | | |
|---|---|---|---|---|---|
| C2808T (8) | - | - | **C2229T** | - | |
| C2814T (8) | - | - | C2259T (6) | - | |
| A2838G (7) | - | - | **C2271T** | - | |
| T2868C (7) | - | - | T2292C (6) | - | |
| T2889C (8) | - | - | A2295G (6) | - | |
| **G2911A** | E987K | 0 | **G2307A** | - | |
| A2913C (9) | E987D | 3 | **T2320C** | S790P | 1 |
| **A2933G** (2) | K994R | 3 | **G2325A** | W791STOP | |
| A2976G (9) | - | - | G2328A (6) | - | |
| T2988A (10) | - | - | T2340C (6) | - | |
| C3030T (7) | - | - | **C2345T** | A798V | 0 |
| A3033G (7) | - | - | G2349A (5) | - | |
| C3057T (7) | - | - | **G2359A** | V803I | 4 |
| T3075C (8) | - | - | **G2364A** | - | |
| G3084A (8) | - | - | **C2376T** | - | |
| A3087G (8) | - | - | A2382G (4) | - | |
| G3093A (8) | - | - | **G2385A** | - | |
| C3096T (6) | - | - | T2388C (3) | - | |
| | | | **G2403A** | - | |
| | | | **G2405A** | C818Y | 0 |
| | | | **G2416A** | G822R | -3 |
| | | | G2418A (5) | - | |
| | | | **G2424A** | - | |
| | | | C2427T (4) | - | |
| | | | **C2439T** | - | |
| | | | **G2457A** | - | |
| | | | C2466T (4) | - | |
| | | | T2478C (5) | - | |
| | | | **C2489T** | A846V | 0 |
| | | | T2502C (5) | - | |
| | | | C2514T (5) | - | |
| | | | G2531C (5) | S860T | 1 |
| | | | **C2533T** | - | |
| | | | C2541T (6) | - | |
| | | | **C2543T** | T864I | 0 |
| | | | A2544T (6) | - | |
| | | | **G2554A** | G868S | 1 |
| | | | T2575C (8) | - | |
| | | | **G2578A** | G876R | -3 |
| | | | **C2592T** | - | |

| | | | | | |
|---|---|---|---|---|---|
| | | | **T2597C** | F882S | -3 |
| | | | **C2600T** | T883I | 0 |
| | | | **C2604T** | - | |
| | | | **G2614A** | V888I | 4 |
| | | | **C2626T** (2) | - | |
| | | | **G2628A** | - | |
| | | | **C2633T** | A894V | 0 |
| | | | T2640C (8) | - | |
| | | | T2655C (8) | - | |
| | | | **C2663T** | S904F | -3 |
| | | | T2664C (8) | - | |
| | | | **C2693T** | A914V | 0 |
| | | | **C2708T** | P919L | -3 |
| | | | **C2714T** | S921L | -3 |
| | | | **C2723T** | P924L | -3 |
| | | | **G2727A** | - | |
| | | | **A2730G** | - | |
| | | | **G2731A** | G927A | 1 |
| | | | **C2736T** | - | |
| | | | **G2743A** | E931K | 0 |
| | | | T2754C (6) | - | |
| | | | G2763A (6) | - | |
| | | | C2766T (6) | - | |
| | | | C2808T (6) | - | |
| | | | C2814T (6) | - | |
| | | | **G2829A** | M959I | 2 |
| | | | A2838G (7) | - | |
| | | | **G2844A** | - | |
| | | | **C2847T** | - | |
| | | | **C2849T** | T966I | 0 |
| | | | T2868C (7) | - | |
| | | | **G2872A** | V974I | 4 |
| | | | **T2879C** | F976S | -3 |
| | | | **G2881A** (2) | E977K | 0 |
| | | | **G2886A** | - | |
| | | | **G2888A** | G979D | 1 |
| | | | T2889C (7) | - | |
| | | | **G2906A** | R985K | 3 |
| | | | **G2907A** | - | |
| | | | **A2908G** | N986D | 2 |

| | | | | | |
|---|---|---|---|---|---|
| | | | A2913C (7) | E987D | 3 |
| | | | **C2918A** | T989N | 0 |
| | | | **C2924T** | A991V | 0 |
| | | | **G2935A** | G995R | -3 |
| | | | **G2966A** | G1005D | 1 |
| | | | A2976G (5) | - | |
| | | | **T2988C** | - | |
| | | | **C2989T** | - | |
| | | | T2988A (5) | - | |
| | | | **G2998A** | D1016N | 2 |
| | | | **G3021A** | - | |
| | | | C3030T (5) | - | |
| | | | A3033G (5) | - | |
| | | | **G3038A** | C1029Y | 0 |
| | | | **A3054G** | - | |
| | | | C3057T (5) | - | |
| | | | T3075C (4) | - | |
| | | | G3084A (4) | - | |
| | | | A3087G (4) | - | |
| | | | G3093A (4) | - | |
| | | | C3096T (4) | - | |
| | | | **G3098A** | C1049Y | 0 |
| | | | **G3119A** | G1056E | 0 |

[a]Residue numbering is according to RVFV strain SA-75, accession number DQ380189. The number of clones in which the mutation is found is given in parenthesis. Bold-face indicates mutations found only in absence (first column) or presence (fourth column) of favipiravir.

[b]PAM250 is the amino acid substitution frequency score in which -7 represents minimum acceptability and +7 maximum acceptability, relative to random sequences (1)
